# TGH: Trans-Generational Hormesis and the Inheritance of Aging Resistance

**DOI:** 10.1101/127951

**Authors:** Ng Li Fang, Jahnavi Suresh, Krishna Chaithanya, Xiao Linfan, Tuan Zea Tan, Jan Gruber, Nicholas S. Tolwinski

**Author notes:** Block MD6, Centre for Translational Medicine, NUS Yong Loo Lin School of Medicine, 14 Medical Drive, Level 10 South, 10-02M Singapore, 117599 (65) 6601 3092.

## Abstract

Animals respond to dietary changes by adapting their metabolism to available nutrients through insulin and insulin-like growth factor signalling. Restricting calorie intake generally extends life and health span, but *Drosophila* fed non-ideal sugars such as galactose are stressed and have shorter life spans. Here, we report that although these flies have shorter life spans, their offspring show significant life extension if switched to a normal sugar (glucose) diet. We define this as TGH or trans-generational hormesis, a beneficial effect that comes from a mild stress. We trace the effects to changes in stress responses in parents, ROS production, effects on lipid metabolism, and changes in chromatin and gene expression. We find that this mechanism is similar to what happens to the long lived *Indy* mutants on normal food, but surprisingly find that *Indy* is required for life span extension for galactose fed flies. *Indy* mutant flies grown on galactose do not live longer as do their siblings grown on glucose, rather overexpression of Indy rescues lifespan for galactose reared flies. We define a process where sugar metabolism can generate epigenetic changes that are inherited by offspring, providing a mechanism for how transgenerational nutrient sensitivities are passed on.

## Introduction and Results

Almost a century ago, Osborne observed plasticity in aging by finding that rats subjected to caloric restriction (CR), experienced delayed and restricted growth, but also lived longer^1^. In the 1930s, McCay continued this work by investigating the link between CR, developmental trade-offs and lifespan ^2^. Now, a century later, CR is recognized as a major, evolutionarily conserved, longevity intervention that extends lifespan, increases stress resistance, and delays the onset of age-associated disorders in organisms as diverse as yeast, flies, rodents and monkeys ^3-7^. The exact mechanism of CR remains only partially understood, but the benefits of CR are mediated by evolutionarily conserved nutrient sensing signalling pathways that trigger increased investment in somatic maintenance at the expense of reproduction and stress response ^8^. These pathways, therefore, not only control the beneficial effects of CR but also the trade-offs associated with these benefits, such as reduced growth, delayed development, and reduced fertility ^8^. For example, women experiencing significant CR stop menstruating, and similarly mouse litter sizes decrease dramatically ^9^. If nutrient shortages are encountered during pregnancy, some nutritional stresses can also be passed on to the next generation ^10^. For many years, information was not thought to flow from soma to germline, but recent studies have shown that offspring are changed epigenetically by environmental factors affecting parents ^11-18^. For example, studies of the link between nutrition and epigenetic inheritance have found that maternal CR can result in metabolic disease in offspring ^19^. Barker at al. suggested that this can be explained by the “thrifty genotype” hypothesis, where foetal epigenetic reprogramming leads to extreme efficiency in nutrient utilization and fat storage ^19^. This hypothesis explains that the detrimental health outcomes are due to a mismatch between the epigenetically expected and actually encountered environment. Evolutionarily, this transgenerational effect can be viewed as a continuum with CR ^20^. Faced with severe lack of nutrients it makes evolutionary sense for the parent to favour somatic maintenance over proliferation essentially waiting in the hope that conditions for reproduction improve. If such shortages occur during gestation, a thrifty phenotype allows the reprogramming of the foetus to follow an appropriate developmental trajectory, thereby increasing the chance of survival for the offspring in a likely nutrient poor environment. For human health, there is great interest in CR and the specifics of epigenetic reprograming for disease risk. In particular, CR mimetic drugs could be used as treatments for human disease and possibly for extending life ^21,22^.

### Transgenerational lifespan effects

We investigated transgenerational effects of a very specific type of dietary modification, where flies were grown on food containing the non-optimal sugar galactose (GAL) as the primary sugar source and compared to flies grown on food containing the basic Kreb’s cycle sugar glucose (GLU) ^23^. CR extends the lifespans of both *Drosophila* and *C. elegans*, but changing the main sugar source from GLU to GAL reduces fly lifespans ^24^. The suggestion is that metabolism of GLU may increase ROS mediated damage and thereby shorten the lifespan of flies ^24^. Expanding on this finding, the thrifty phenotype hypothesis suggests an epigenetic mechanism for transgenerational reprogramming of the genome for a given environment. We therefore investigated how detrimental effects in the parental generation affect offspring. In particular, we asked if a mismatch between the expected and the actual environment can be beneficial rather than, as with the thrifty phenotype, detrimental.

In our experiments, there is a small reduction in *Drosophila* lifespan when reared on GAL as compared to their siblings reared on GLU (Fig. 1A-C). Next, we looked at flies reared on GAL and GLU, but switched to the opposite sugar upon eclosing (Fig. 1D, Schema of the experiment). We found a significant increase in the lifespan of flies reared on GAL, but switched to GLU, especially when compared to flies reared on GLU and switched to GAL (Fig. 1C, please note the extension in total lifespan and average lifespan). In other words, flies whose genome “expected” a more hostile environment than actually encountered showed increased lifespan while in the opposite case lifespan was shortened. To ensure that this effect did not stress the flies into inactivity, we looked at the fecundity of flies reared on GAL. We observed that peak fecundity was delayed by one or two days in GAL flies, but overall offspring numbers did not differ significantly (Fig. 1E-F, Table).

**Figure 1:**
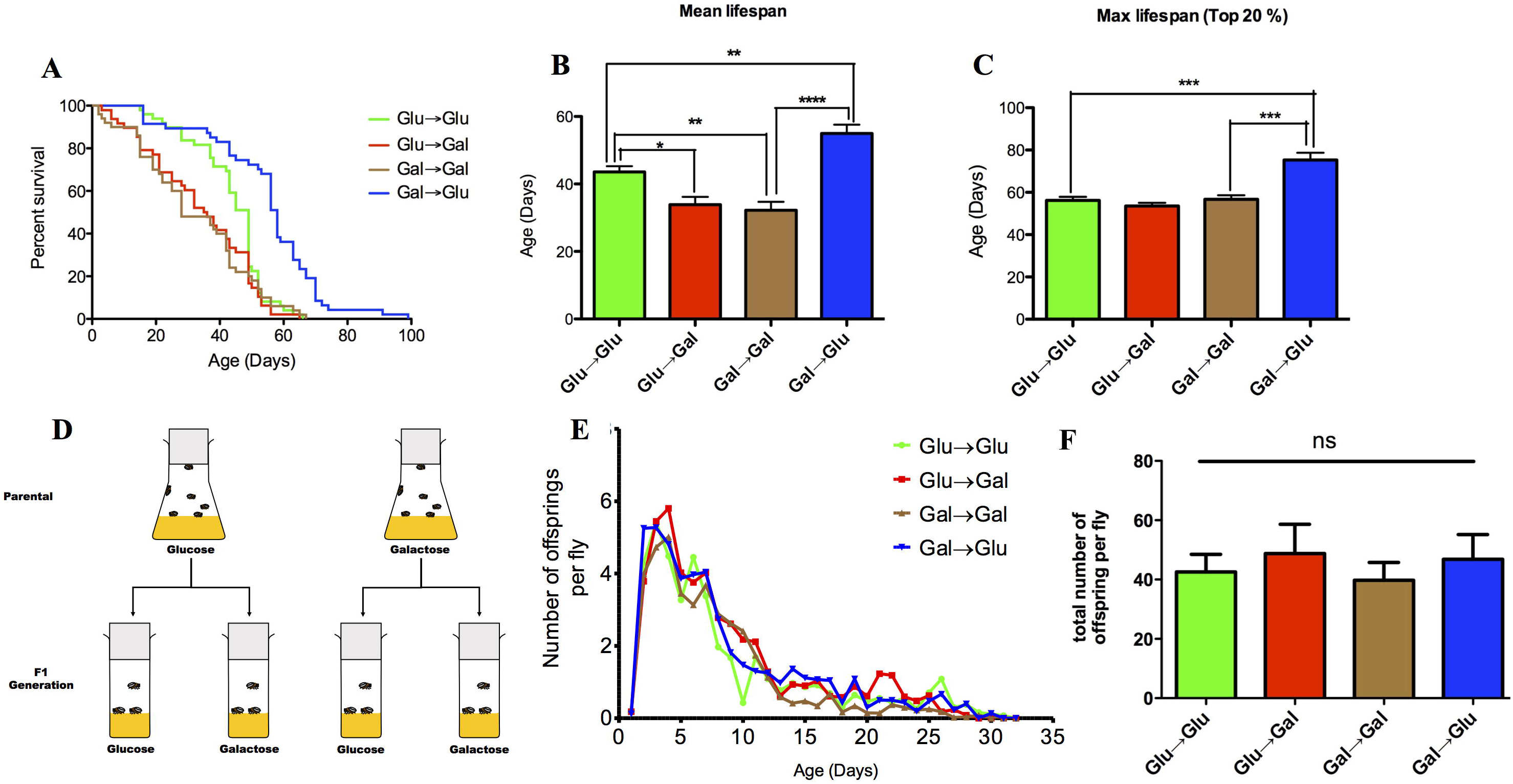
Survival and fecundity of Drosophila grown on glucose and galactose (n=50 flies per condition per study). (**A**) F1 generation of flies shifted from Gal → Glu had significant lifespan extension, nearly twice as long as F1 generation flies fed from Glu→ Glu (p<0.05, log-rank (Mantel-Cox) test). (**B**) Flies shifted from Gal→Glu showed significant mean lifespan (26%) extension compared to Glu→Glu flies (p<0.01). Flies switched Glu→Gal and Gal→Gal show significantly shortened mean lifespan by 22 and 26 %, respectively (p<0.05 and p<0.01, respectively). (**C**) A significant maximum lifespan extension of 34 % was observed in Gal→Glu flies relative to Glu→Glu control flies (p<0.001). Mean and maximum lifespan were analyzed using ANOVA with Bonferroni’s post-test, mean±SEM. (**D**) Schema of the experiment: Parental generation of flies was raised on glucose and galactose. F1 generation of flies from both Glu and Gal were fed with both Glu and Gal food. (**E**) A typical reproductive schedule and cumulative offspring produced over a lifetime for flies exposed to glucose and galactose. Average number of offspring produced was quantified for a period of 32 days. Points represents number of offspring per fly, mean±SEM. Flies raised from Gal→Glu reproduced earlier than flies fed from Glu Glu, Glu Gal and Gal Gal. (**F**) Flies switched from Gal Glu appeared to produce more progeny than Glu→Glu flies over their total lifetime fecundity. Gal→Glu compared to Gal→Gal showed a trend towards lower egg laying capacity, but these trends were not statistically significant.

**Table 1.** Peak of fertility for flies exposed to Glu and Gal. Gal→Glu flies produced the most offspring throughout their lifetime with a peak on day 2, earlier than Glu→Glu, Glu→Gal and Gal→Gal. During the peak of fertility, the mean number of offspring for Gal→Glu flies was 5.25±0.80.

| Treatment | Peak of Fertility |  |
| --- | --- | --- |
|  | Peak Time (Day) | Number of offspring<br>± SEM (fly) |
| Glu→Glu | 3 | 5.42±0.81 |
| Glu→Gal | 4 | 5.80±0.69 |
| Gal→Gal | 4 | 5.01±0.57 |
| Gal→Glu | 2 | 5.25±0.80 |

The Barker hypothesis states that organisms that suffer a cycle of starvation can pass along to their offspring an expectation of poor diet in their future ^19^. This epigenetic process leads to positive effects if the offspring do encounter the expected environment, but can also be detrimental if the environment differs significantly. When applied to calorie or nutrient availability the hypothesis explains why foetal starvation can lead to diabetes and obesity in a generation that encounters a nutrient rich environment that their parents didn’t. GAL can should be a poorer energy source compared to GLU, and we had expected that GAL to GLU switched flies would suffer detrimental lifespan and health effects, but we observe the opposite effect or the extension of fly lifespan. Flies from GAL parents that encounter GLU as the primary sugar source live longer. However, in addition to being more difficult to metabolise, metabolism of GAL can also stress flies ^24^, so we looked at mitochondrial ROS in GAL flies and confirmed that mitochondrial ROS was indeed elevated (MitoSOX, Fig. 2A). We confirmed this result using the less specific global ROS dye DCF-DA (Fig. 2B). To further support the notion that the metabolism of GAL imposes oxidative stress, we also measured oxidative modification to protein as measured by the protein carbonyl (PCC) assay (Fig. 2B). Again, we observe a consistent increase in PCC in GAL reared flies as compared to GLU control flies regardless of genetic background (Fig. 2C). These data show that flies reared on GAL are exposed to higher levels of oxidative stress than those reared on GLU.

**Figure 2:**
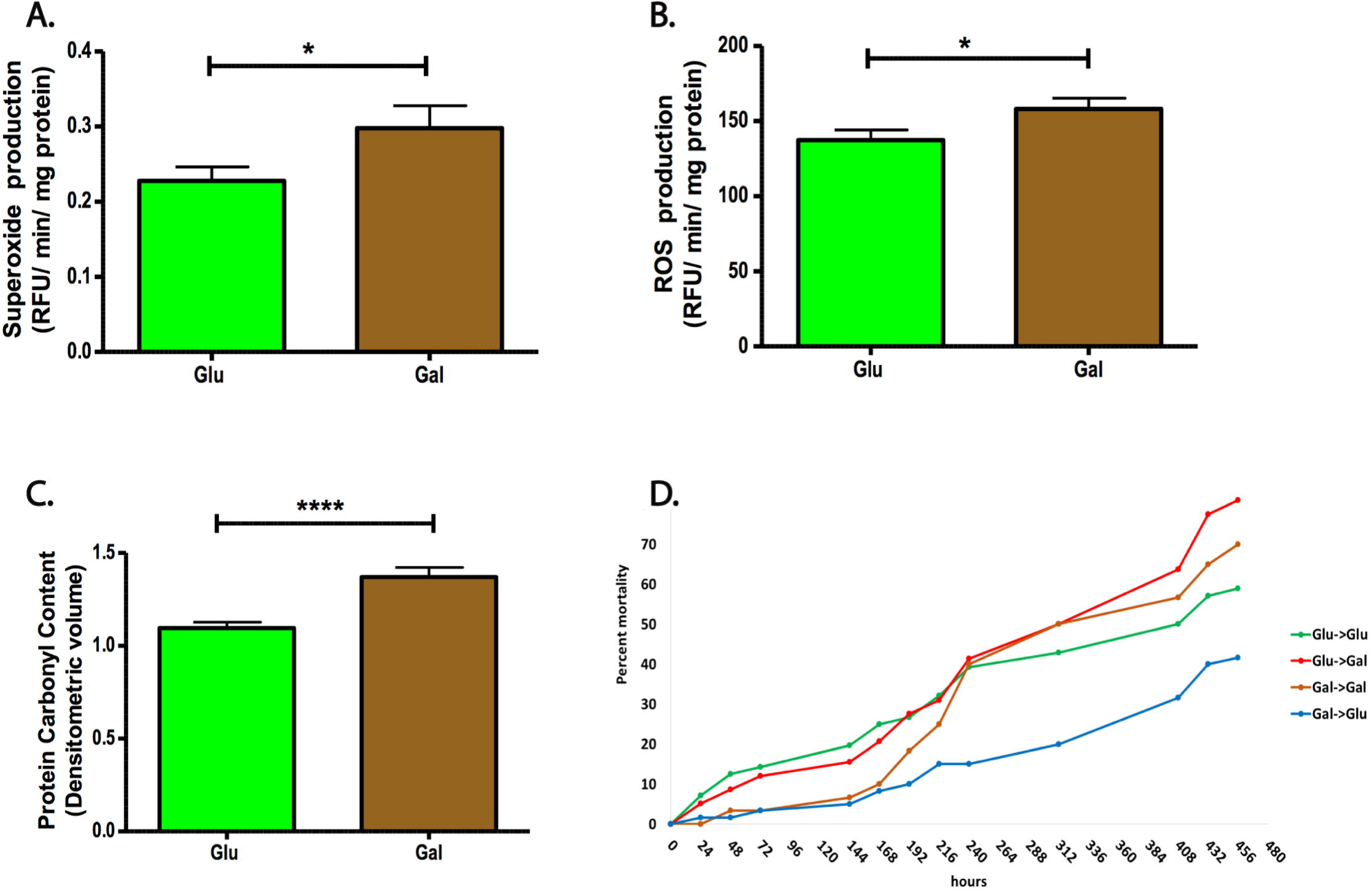
Effect of Glucose and Galactose exposure on flies’ resistance to exogenous stress and global oxidative stress. (**A**)MitoSOX Red fluorescence assay as an indicator of mitochondrial superoxide generation from six independent experiments. Significant elevation of MitoSOX red fluorescence level was detected in Gal fed flies compared to Glu fed flies (p<0.05, Student’s t-test, mean ±SEM), n=18 independent replicates. (**B**) DCF-DA fluorescence assay from six independent experiments. Flies fed with Gal had a significantly higher global ROS than flies fed with Glu (p<0.05, Student’s t-test, mean ±SEM), n=18 independent replicates. (**C**) Global oxidative stress. Relative protein carbonyl content normalized to protein from three independent experiments, measured by a densitometric analysis of the spot-blot intensity. Gal fed flies had significantly higher protein carbonyl content compared to Glu fed flies (p<0.0001, Student’s t-test, mean ±SEM), n=9 independent replicates. (**D**) Mortality curve of flies treated with paraquat. Parental generation of flies fed with Glu are less resistant to paraquat in the first 9 days of treatment. Subsequently, after day 9, flies fed from Gal→Glu are more resistance to paraquat. The acute treatment of Gal→ Glu flies with paraquat induced only 20% mortality after 13 days of treatment whereas Glu→Gal, Gal→ Gal flies treated with paraquat both induced 50 % of mortality and Glu→ Glu flies induced 42.9 % of mortality on the same time point.

This data suggested that in analogy to food scarcity leading to a “thrifty” phenotype, flies switched from GAL to GLU might be epigenetically reprogrammed to exhibit a stress resistant phenotype and would therefore be expected to deal with elevated ROS levels better. We confirmed this expectation by exposing flies to the ROS generating poison paraquat, showing that flies switched from GAL to GLU were far more resistant to oxidative challenge than all other groups (Fig. 2D). This suggests that these flies might be conditioned or epigenetically programmed to survive oxidative stress levels higher than they actually encounter. Similar results regarding elevated stress resistance to paraquat have been shown previously for animals exposed to CR or pre-conditioned with ROS. As for CR, in the absence of the expected stressor (GAL) this elevated stress resistance translates into a longer lifespan.

Barker hypothesis predicts adaptive *epigenetic* changes resulting in transgenerational transcriptional changes tailored to the environmental conditions expected based on environmental conditions during pregnancy. We observed transgenerational adaptation where the F1 generations were fully adapted to surviving on their respective sugar (Fig. 1D). The short time in which this adaptation appeared was insufficient for any adaptive genetic mutations to have propagated throughout the population suggesting an epigenetic explanation. We therefore began with transcriptome sequencing of flies from the two sugar conditions. RNA sequencing from adults reared on the two sugars showed 484 reads of 11496 as altered (Fig. 3A, by FPKM). We further refined these changes to 87 genes changing at significant rates between the two conditions with 75 upregulated and 12 repressed in GAL reared flies compared to GLU (Fig. 3B). We reasoned that changes in chromatin modifications would be the likely basis of RNA expression changes so we proceeded to perform Chromatin Immunoprecipitations with H3K27me3 antibody followed by sequencing (ChIP-Seq). As this is a transcriptionally repressive chromatin mark, we used peaks in ChIP-Seq data correlated with RNA-Seq data to find genes that showed higher expression and lower H3K27me3 signal, and vice versa (Fig. 3C). We observed 38 genes that showed this pattern (Fig. 3C), further refining the gene list from 87. We grouped these genes by gene ontology (GO) terms and found an expected enrichment in metabolic and signal transduction genes (Fig. 3D). An example of the data is shown (Fig. 3E) where expansion of H3K27me3 signal in GLU flies correlates with a lower RNA read number in the sequencing data. We further analysed the ChIP-Seq data by confirming that sequencing data was corroborated by ChIP-qPCR by testing 10 random genes (Not Shown). We also looked at the distribution of H3K27me3 peaks around genes, and found that the biggest difference was observed in promoter regions with GLU reared flies having nearly twice as many promoters methylated (Fig. 3F), suggesting that a lot of genes are activated when flies are reared on GAL a finding corroborated by the increase in gene transcription in these flies (Fig. 3B). We classified these genes according to GO terms finding little different in GLU flies, but observed many changes in general functions such as transcriptional regulation and metabolism, but no change in the many genes previously associated with aging and, in particular, no changes in gene expression of any of the canonical antioxidant genes (Fig. 3G). As these data did not point to an obvious mechanism we used KEG analysis to look for pathways affected by the gene expression changes. We observed four pathways increasing, immune system and protective other than immune system pathways suggesting a stress response, carbohydrate metabolism pathway which fits with the change in sugar diet, and lipid metabolism (Fig. 3H). The first two pathways were interesting but expected, whereas the finding of the lipid metabolism pathway was unexpected.

**Figure 3:**
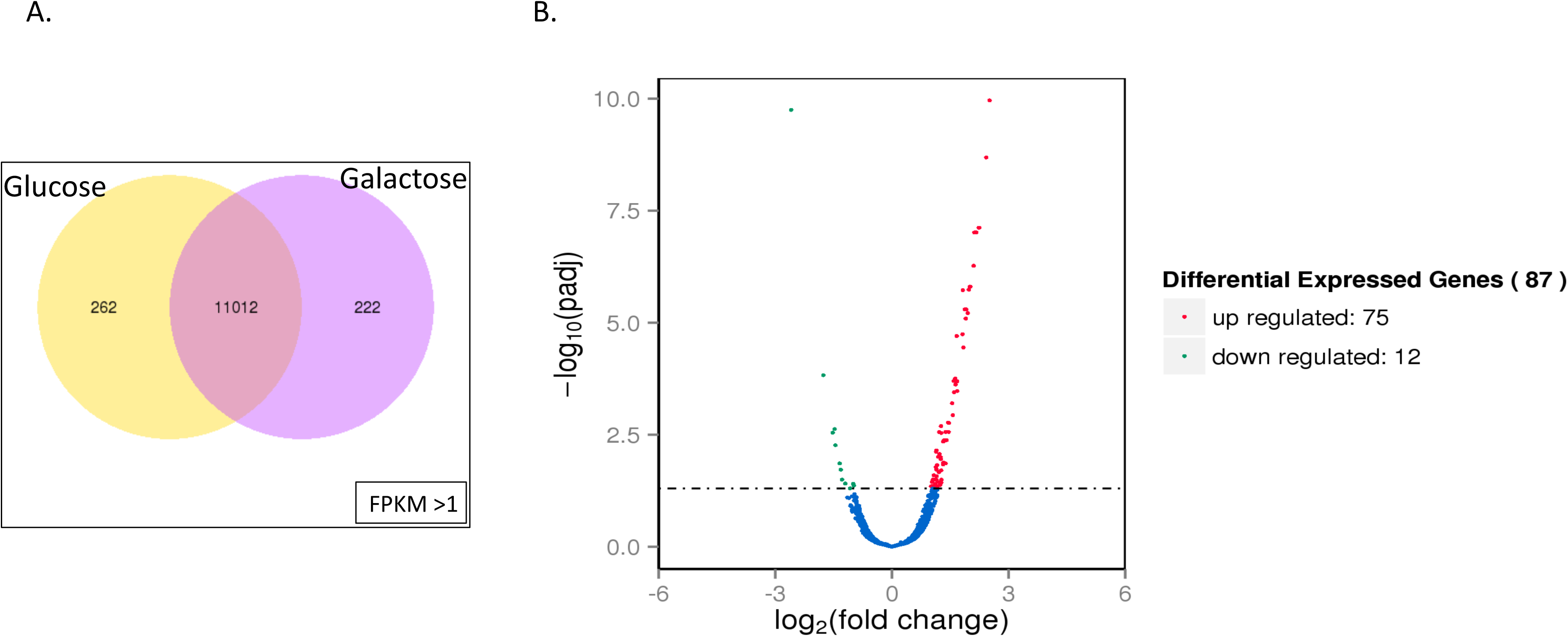
RNA and ChIP Sequencing Analysis: (**A**) Total number of expressed genes from transcriptome sequencing data of 5day old Glu and Gal fed flies. Intersection of the Venn diagram is showing the expressed genes overlapping between the two groups. FPKM>1 represents the expression threshold. (**B**) Volcano plot representing the differentially expressed genes upon analysis of transciptome sequencing data using DEseq2 package. Of the 87 differentially expressed genes, 75 genes were up regulated and 12 genes were down regulated in the Gal fed flies as compared to Glu. (**C**) Heat map represents the gene expression coverage data of transcriptome and the corresponding H3K27Me3 repressive chromatin peak signal from ChIP sequencing data. Heat maps represent the overall higher expression levels of the genes in the transcriptome of 5 day old Glu fed flies and the corresponding lesser repressive chromatin signal from the ChIP sequencing data in these flies. Comparatively high chromatin methylation is found in the genome of Gal fed flies. (**D**) Figure showing the gene ontology enrichment data of Gal fed flies. In support of the higher repression signal from the ChIP sequencing data, enrichment of metabolic processes is found in Gal fed flies, but no significant gene ontology processes were found from the glucose fed flies. (**E**) Snap shot from integrative genomic viewer which represents the gene expression coverage peaks from transcriptome and ChIP sequencing data where the expression levels are in the opposite direction. (**F**) Bar diagram represents the distribution of repressive chromatin peaks in different parts of the *Drosophila* genome. Peak distribution data from both glucose and galactose fed flies have been represented (5day old flies). Repressive chromatin peaks are more in the promoter region of Glucose fed flies compared to galactose fed flies. (**G**) Gene ontology enrichment of those genes having repressive chromatin peaks in the promoter regions of both glucose and galactose fed flies. Enrichment of metabolic and biosynthetic processes is high in galactose fed flies but not in the case of glucose fed flies. (**H**) KEG analysis showing four processes enriched.

In agreement with the Barker hypothesis, we observe epigenetic changes that result in adaptive gene expression changes in flies reared on different sugars. Changes in lipid metabolism could explain the lifespan extension as recent work links lipids to longevity determination rather than simply as required structural components of cell membranes and energy storage molecules ^25,26^. The level of n-3 polyunsaturated fatty acids (PUFAs), in particular, bound to membrane phospholipids has been suggested as a key determinant of lifespan and aging ^27,28^. The key is the ratio between monounsaturated fatty acids (MUFAs) and PUFAs. The degree of unsaturation determines the susceptibility of fatty acids to oxidative damage with less saturated lipids being prone to oxidation leading to oxidative damage across the membrane (free radical chain reactions)^27,29^. A higher MUFA to PUFA ratio therefore is associated with increased resistance to oxidative damage. Indeed, genetic and dietary modification associated with lifespan extension have been shown to increase MUFA:PUFA in *C. elegans* and mice ^25,30^. Elevated MUFA:PUFA ratios in plasma membranes have even been suggested to be related to lifespan in humans ^29^. Consistent with these previous findings, we observed an elevated MUFA:PUFA ration in GAL flies (Fig. 4A-B). To examine this further, we turned to overall lipid composition or a lipidomics approach to discover whether lipid-related gene expression changes correlate with changes in the lipid composition in flies. We found that a number of aging-protective phospholipids were present at higher concentrations in flies reared on GAL as compared to those reared on GLU (Fig. 4C). For example, phosphatidyethanolamine (PE), which when reduced in yeast leads to an acceleration in aging-associated production of ROS and finally death while overexpression extended lifespan ^31^, showed an increase of 33% on Day 1 reaching 44% on Day 5 in GAL flies (Fig. 4C). Next, we looked at sphingomyelins (SM), lipid molecules that share a similar head group to phosphatidylcholine (PC) but with a ceramide backbone. SMs are enriched in plasma from long-lived individuals, and levels of its breakdown product ‘ceramide’ a pro-aging signalling molecule are reduced ^29^. We observed an increase in the total composition of SM an 8% increase on Day 1 and a 11% increase on Day 5 (Fig. 4D), and a drop in ceramide levels 7% Day 1 and a 14% Day 5 in GAL flies (Fig. 4D). Another aging-protective lipid PC ^32^ showed an increase of 22% on Day 1 and 19 % increase on Day 5 (Fig. 4D). We also observed a 25% decrease in triacylglycerides (TAG) on Day 1 and 9 % at Day 5 (Fig. 4D). TAG accumulation is linked to aging through a number of factors ^33,34^, and mice with lower concentrations of TAGs live longer ^35^. These studies suggest that epigenetic changes changing the expression of lipid related genes affected lipid composition of the membranes on flies and correlated with lifespan extending changes in lipid composition.

**Figure 4:**
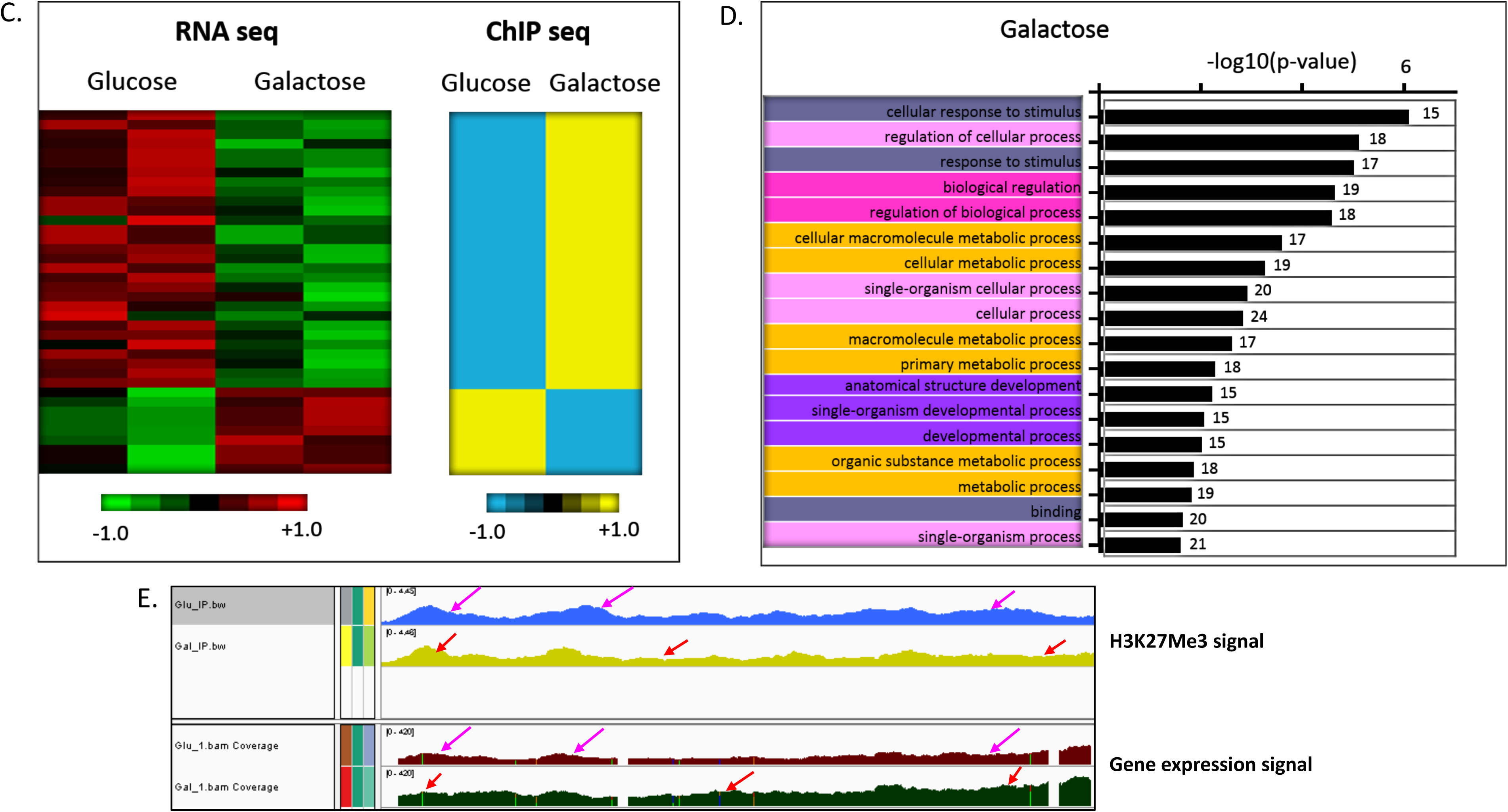
Lipidomic Analysis of Drosophila shows an increase in life span protective membrane lipids in Gal reared flies. (**A**) MUFA/PUFA ratio showing an increase in the phosphatidylcholines containing monounsaturated (MUFA) vs. polyunsaturated fatty acid (PUFA) ratio in Gal flies. The MUFA data shown here represents fatty acids containing one double bond while the PUFAs data point is a representation of all the pooled fatty acids containing double bonds ranging from 2 to 5 double bonds. Error bars indicate standard deviation. (**B**) MUFA/PUFA ratio increases over time as shown by 1 day old vs. 5 day old Gal treated flies. (**C**) Multipanel pie charts showing the effect of diet on lipid class abundance in *Drosophila*. The amount of each lipid class in Drosophila reared on Glu and Gal on Day 1 and Day 5. Lipid class amount is calculated as molar % with respect to total lipids. Lipids quantified include phosphatidylcholine, phosphatidylethanolamine, ceramides, sphingomyelin, DAG and TAG. Dramatic increases in PE and PC and significant decrease in TAGs are observed. (**D**) Dot plot representing changes in various lipid classes on Day 1 and Day 5 in Gal treated flies relative to Glu treated flies.

### The Indy Connection

Sugar metabolism is associated with energy production, mitochondria and longevity through regulation of glucose metabolism in a variety of organisms ^36,37^. As our lipidomic approach only looked at total membrane and nonmembrane lipids we could not tell whether mitochondria were affected specifically. To narrow the possible mechanisms, we looked at mutations that affect metabolism and lifespan. One of the best-known aging-protective mutations is known as *I’m Not Dead Yet* or *Indy*^38^. We tested the effect of mutating *Indy* in flies reared on GAL and GLU. As expected, *Indy* mutations extended the lifespan of flies grown on GLU, but surprisingly we found that the same loss of *Indy* was detrimental to flies living on GAL (Fig. 5A-C). Even more remarkable was that overexpression of Indy extended the lifespan of flies grown of GAL but not of flies grown on GLU. The Indy protein is as a plasma membrane cotransporter similar to mammalian cotransporters involved in uptake of di- and tricarboxylic intermediates from the tricarboxylic acid cycle ^39^. *Indy* mutants have longer lifespans by a mechanism similar to CR, including altered lipid metabolism and higher mitochondrial biogenesis ^39^. We looked at the lipid composition of *Indy* mutant and *Indy* overexpression flies by the same lipidomic approach used above. We observed that long-lived flies, *Indy* overexpressing reared on GAL, showed an increase in aging protective PE and a reduction in the poor aging TGC (Fig. 5). The multipanel (Fig. 5D and 5E) shows lipid class specific analyses in *Indy* mutants, both loss of function *Indy^302^* and overexpression (UAS-Indy). We found significant differences across all lipid classes, for instance, *Indy^302^* on GAL show a 50% decrease in TAG levels on both Day 1 and 5, while the drop was 30% when reared on a GLU diet. TAG levels dropped by 10% in UAS-Indy flies on GAL food on Day 1 and a 40% drop on Day 5. Both *Indy^302^* and UAS-Indy showed similar DAG profiles on GLU.

**Figure 5:**
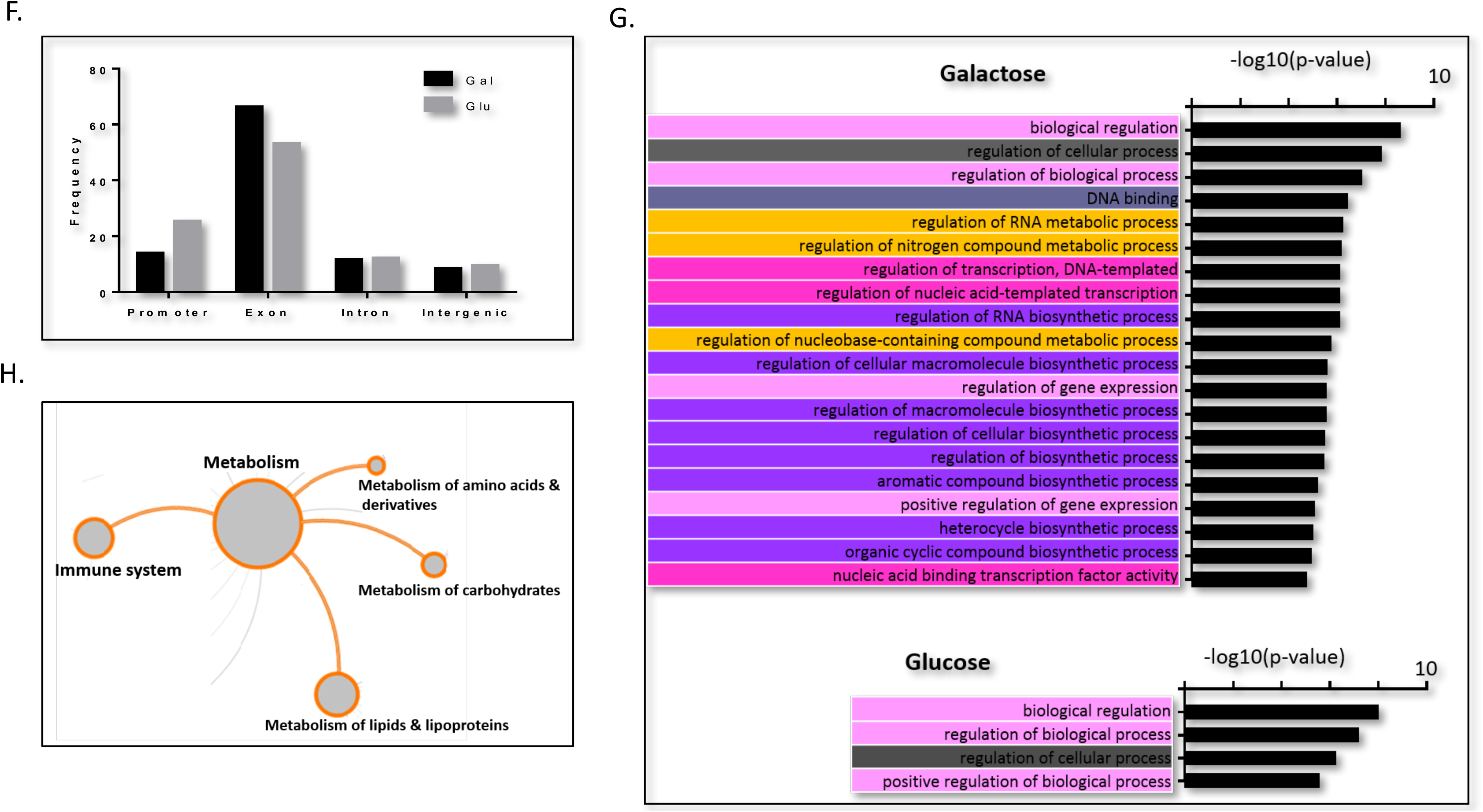
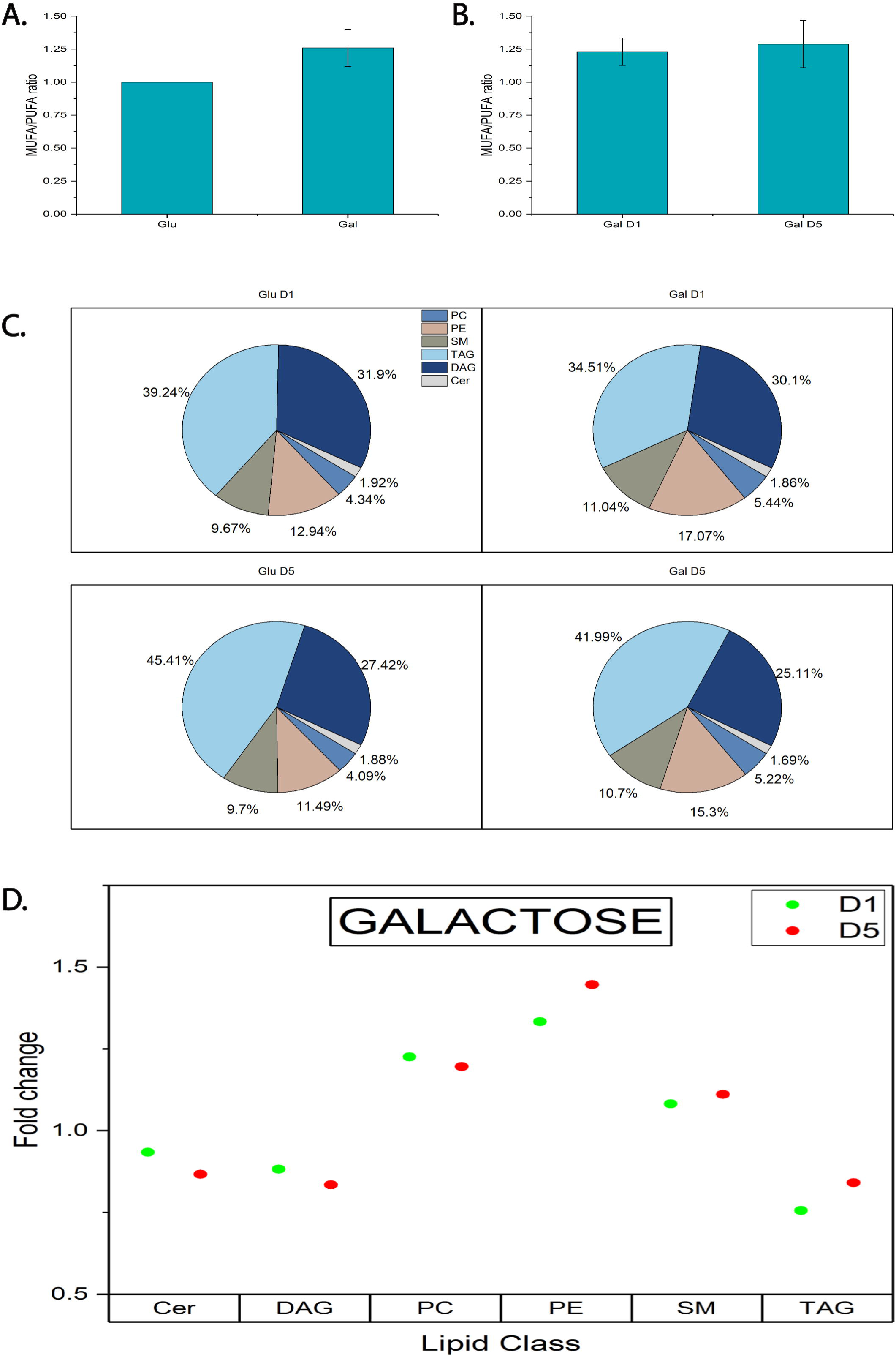
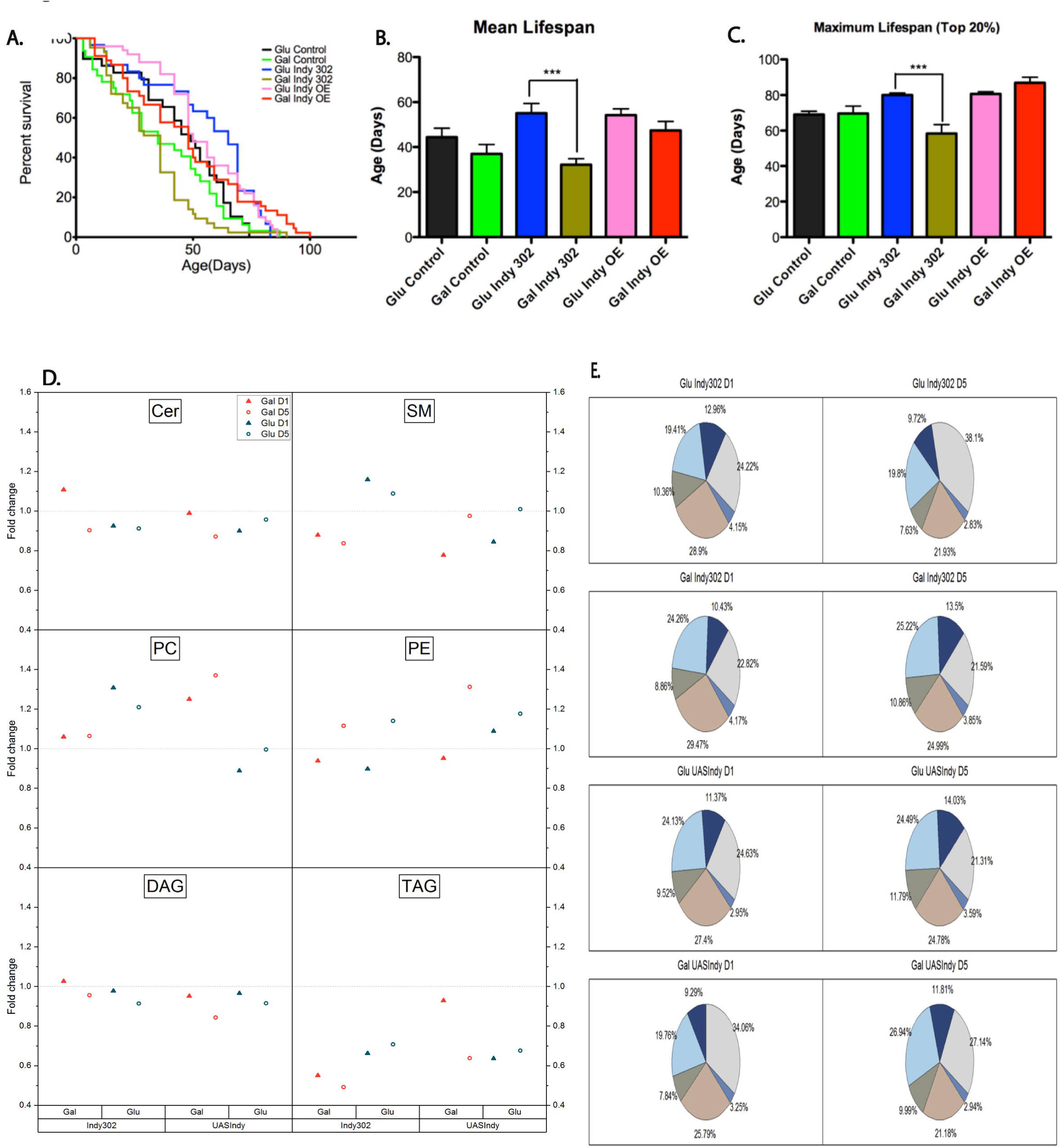
Lifespan extension in Indy flies. (**A**) *Indy^302^* flies reared on Gal had significantly shorter lifespan that *Indy^302^* flies fed with glucose (p<0.0001, log-rank (Mantel-Cox) test). (**B**) *Indy^302^* flies reared on Glu had 24% mean lifespan extension compared to wild type flies reared on Gal. Mean lifespan of *Indy^302^* flies fed with Gal was significantly shortened by 42 % relative to *Indy^302^* flies on Glu. *Indy* overexpression flies fed with Gal had no significant difference in the mean lifespan compared to Indy overexpression flies in Glu. (**C**) Gal supplementation on *Indy^302^* flies significantly shortened the maximum lifespan compared to Glu supplemented *Indy^302^* flies. *Indy* overexpression flies fed with Gal showed a trend towards maximum lifespan extension by 8% relative to flies fed with Gal. Mean and maximum lifespan were analyzed using ANOVA with Bonferroni’s post-test, mean ±SEM. (**D**) Dot plot representing changes in various lipid classes on Day 1 and Day 5 in Gal treated flies relative to Glu treated flies comparing *Indy^302^* to *Indy* overexpression. (**E**) Multi-panel pie charts showing the effect of diet and comparing *Indy^302^* to *Indy* overexpression on lipid class abundance in *Drosophila*. Lipid class amount is calculated as molar % with respect to total lipids. Lipids quantified include phosphatidylcholine, phosphatidylethanolamine, ceramides, sphingomyelin, DAG and TAG. Dramatic increases in PE and PC and significant decrease in TAGs are observed.

Next, we looked at the aging-protective phospholipids, PC and PE. No significant alterations were observed on *Indy*^*302*^ on GAL diet nor UAS-Indy on GLU whereas PC profiles in *Indy*^*302*^ on GLU and UAS-Indy on GAL showed significant increases on 20 – 30 % on both days. These results agree with the hypothesis that PC concentration increases in long lived mutants. Alterations in PE compositions were mainly observed on Day 5 for all the conditions (about a 10% increase) except that UAS Indy when raised on GAL displayed a more than 20% increase in its levels. No major changes in the levels of the pro-aging molecule, ceramide was observed in all conditions except for *Indy^302^* on GAL where a 10% was seen. SM analysis revealed an increase in its levels only in *Indy^302^* reared on GLU while on GAL a more than 10% decrease was observed (Fig. 5). UAS Indy flies on both substrates showed similar trends with almost a 20% decrease on Day 1 only and a return to normal levels on Day 5. Taken together, these data suggest that GAL flies have cells that alter their lipid composition and no longer respond to lower Indy levels, but are very sensitive to increased Indy. It is likely that limited nutrient uptake, perhaps due to different membrane composition, requires extra Indy protein.

In conclusion, our study shows transgenerational, epigenetic inheritance of an adaptive response to nutrient induced stress. We have termed this TGH, or trans-generational hormesis as we observed an inherited adaptation to elevated oxidative stress that protects offspring. The parents are stressed by living only off GAL, leading to higher ROS levels, oxidative damage and shortened lifespan. This nutritional stress elicits an adaptive response in gene expression that results in pronounced changes in membrane lipid composition, making them more resistant to oxidative damage. These changes are transferred to the next generation through an epigenetic mechanism, regardless of the sugar (GLU/GAL) consumed by the offspring itself. Offspring that do not face the GAL dependent ROS increase are therefore benefiting from lifespan extension by a hormetic response to the stress experienced by their parents. Further, our sugar based model reveals a new protective function of the Indy metabolite co-transporter.

This sugar based life extension assay can be used to study the basic mechanisms of aging as well as linking diabetes to aging and lipid accumulation. This model organisms approach allows for a rapid linking of nutrition to neurodegeneration ^15,40^. There are obvious medical and public health implication of epigenetic inheritance of nutritional metabolic risk ^41^, as several studies have proposed that somatic alterations in diet or other environmental changes can affect offspring ^11-19^. Barker’s thrifty hypothesis states that a thrifty environment during foetal development can modify the offspring to expect a poor environment in the future. One example of this is the Dutch famine at the end of World War II where individuals exposed to the poor environment during gestation had a higher incidence of diabetes than those born the year before. Other neonatal effects have been observed on offspring weight, reviewed in ^10^. These effects are believed to be based on epigenetic changes that can be reversed in vivo, as recent work has shown that aging is a reversible process specifically through epigenetic reprograming ^42^. This process requires transient expression of the Yamanaka factors, but it should be possible to mimic that effect with drugs ^42^. These factors work by resetting the epigenetic state of cells in the body making them more youthful. We show that even a small dietary change in sugar can affect the epigenetic state opening up the possibility that more specific drug based treatments will target epigenetic states and enhancing life and health span.

